# Camera Paths, Modeling, and Image Processing Tools for ArtiaX

**DOI:** 10.1101/2024.09.23.614454

**Authors:** Utz H. Ermel, Deborah Moser, Pauline Roth, Gunnar Arctaedius, Maren Wehrheim, Margot P. Scheffer, Achilleas S. Frangakis

## Abstract

The enhancement of biomolecular image analysis and data interpretation is significantly improved through the application of advanced visualization techniques. Numerous visualization packages are currently available, spanning a broad spectrum of applications. Recently, we have extended the capabilities of UCSF ChimeraX to address the specific demands of cryo-electron tomography. Here, we introduce the evolution of our existing plugin, ArtiaX, designed to generate models that facilitate particle selection, define camera recording paths, and execute particle selection routines. In particular, diverse models can be generated and populated with putative particle positions and orientations. A specifically tailored coarse grained algorithm was developed to rectify overlaps, as encountered in template matching, employing a rapid and efficient approach. In addition, models can be used to drive the camera position, thereby simplifying the process of movie creation. The plugin incorporates fundamental image filtering options for the on-the-fly analysis of tomographic data and also provides compatibility of particle lists with RELION-5 star files. Collectively, this update of ArtiaX comprehensively encompasses essential tools for the analysis and visualization of electron tomograms. It retains its hallmark attributes of speed, reliability, and user-friendliness, fostering seamless human-machine interaction.

## Introduction

Interactive visualization of biomolecular images is an inherent component of data analysis and interpretation. Many tools for automated, machine-learning based annotation have been developed in recent years, yet manual labelling or curation for generation of ground truth labels remains a requirement for most of these algorithms. Numerous extendable software packages for interactive 3D visualization are available for the cryoET space (e.g. Amira^1^, UCSF ChimeraX^2^, Napari^3^, Dragonfly^4^ and others). Previously, we developed an open-source toolbox named ArtiaX^5^ on the platform of ChimeraX^2^. The toolbox enables fluent human-machine interaction and improves the visualization of cryoET-related tasks, exploring the VR capabilities of ChimeraX^6^. In particular, the selection and inspection of putative positions and orientations of particles on 2D slices and within 3D images was enabled. The fluent interaction with the datasets allowed for particle lists to be quickly assessed and curated. Users also benefited from the multitude of features of ChimeraX, such as cross-platform operation, animation capabilities, and 3D image file format compatibility.

In this technical report we present several additions that aim to further improve and optimize ArtiaX, allowing users to perform more steps of the typical cryoET analysis pipeline in ChimeraX. Now, macromolecular complexes and other cellular structures can be modelled using geometric primitives, camera recording paths can be defined in 3D, basic image processing for the analysis of tomographic data is enabled, and tools for editing segmentation meshes are provided. The new version of ArtiaX can be downloaded from the ChimeraX toolshed and source code is available on GitHub.

## Results

We first set out to improve the user experience by enabling the state of the plugin and all associated data in ChimeraX sessions to be saved, thus allowing easy storage, recovery and sharing of large projects. This modification also facilitates creating animations and working on larger curation tasks, which usually require hours of work.

### Generation and refinement of models for selecting particle positions

Selecting positions for sub-tomogram averaging is now predominantly performed automatically, e.g. by template matching or neural networks^7,8^. Often – in particular for challenging datasets or for datasets where the underlying object is not known – results of these methods are manually curated or post-processed after running the particle picking algorithm. Previously, individual positions could be manually curated, and particles could be manually oriented in ArtiaX. Here, we introduce the new capabilities of ArtiaX to generate models that can be populated with putative particle positions (Figure 1). Primitives included in ArtiaX are lines/curves, spheres, 2D surfaces and hulls with varying degree of convexity. All geometric primitives are generated by either fitting 3D functions on or computing hulls around user-or algorithm-generated sets of points. This is useful to generate the initial positions and orientations of cytoskeletal filament subunits (curve primitive)^9^, cell-cell junction proteins (2D surface primitive)^10^, viral capsids (sphere primitives) or plasma membranes (hulls)^11^. The spacing between generated points, the method used for distributing them, as well as their orientation can be adapted according to user-defined parameters.

**Figure 1.**
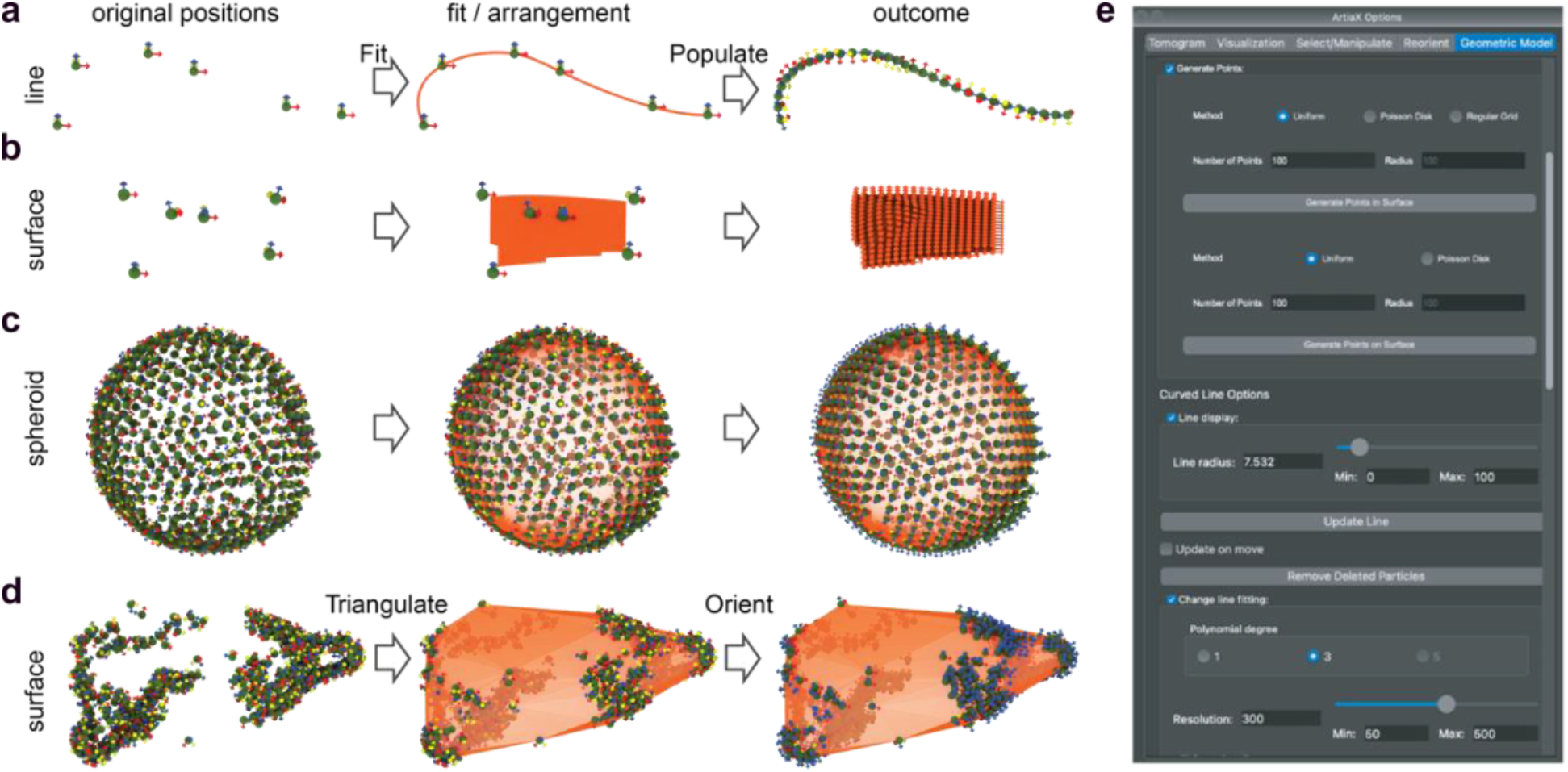
ArtiaX provides geometric primitives for modelling particle positions and shapes. **(a)** Line models (e.g. for modelling filamentous structures such as actin and microtubules) can be created and populated (e.g. evenly spaced) using different - orientation constraints (e.g. helical). **(b)** Surface models can be created from a set of points and further populated with particle positions. **(c, d)** Ultimately complex shapes such as a spheroid object and surfaces of cells can be modelled with an adaptive convexity and particles can be positioned and oriented without requiring dense voxel segmentations. **(e)** The “Geometric Model” tab provides UI elements for manipulating geometric models.

Additionally, segmentations generated using external software can be converted into geometric model objects and used as reference to orient particles, eliminating the need for manual particle orientation.

### Segmentation and Mesh curation tools

Apart from curating and pre-orienting particles, ArtiaX now also facilitates the curation of 3D voxel segmentations and meshes. The plugin provides two new mouse modes, which can be used to remove connected components or patches of triangle meshes and convenient commands and interface elements for converting the curated meshes back to dense voxel segmentations. The same tools can be used to quickly generate segmentations from any of the geometric primitives introduced above. This speeds up generation of ground truth labels for machine learning tasks and curation of segmentations generated using neural networks.

### Verification of position selection and refinement of template matching results

When placing molecules within a tomogram, whether selected through methods such as template matching, neural networks or manual selection, there is a potential for neighbouring positions to overlap. This not only gives rise to an aesthetic concern as overlapping particles become apparent during visualisation, but also contains duplicate particles contributing signal to a sub-tomogram average.

To address this issue, ArtiaX now introduces an algorithm designed to eliminate these overlaps. The algorithm first quantifies the overlap by either calculating the depth of overlap between particles or by using a Monte Carlo method combined with Poisson-Disc sampling to assess the volume overlap between 3D surfaces. Once the overlap is identified, particles are iteratively repositioned to minimize the overlap. This repositioning can be constrained by additional boundary conditions, such as restricting movement to a user-defined surface (Figure 2).

**Figure 2.**
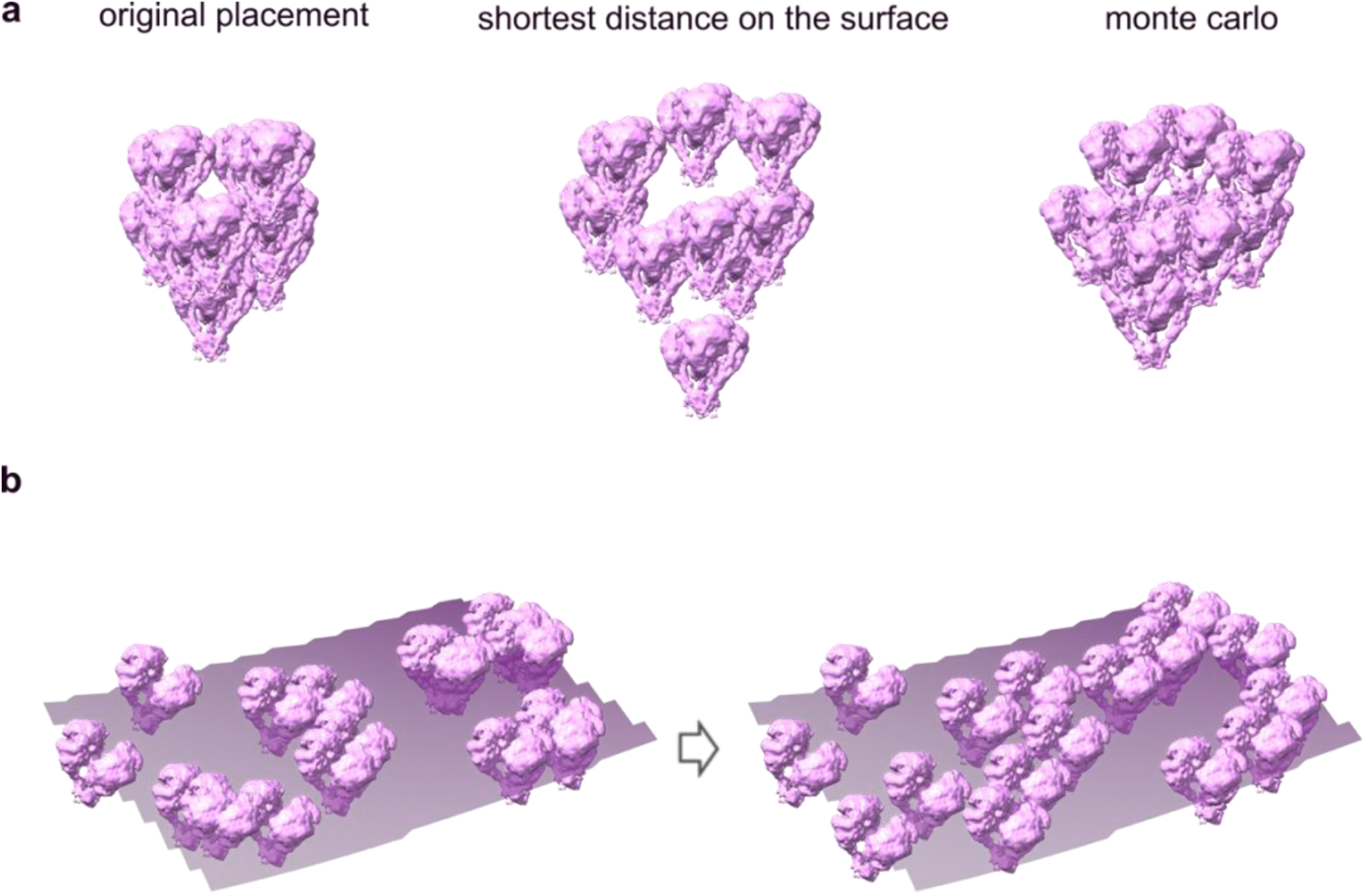
**(a)** Iterative adjustment of objects (EMD-17589) to eliminate overlap, using two different methods to determine the 3D overlap. *Left:* Initial particle positions, *center:* overlap resolved based on the depth of overlap, and *right:* overlap resolved using the Monte Carlo method. **(b)** Overlap removal between particles (EMD-17589) constrained to a user-defined surface.

This method improves the visualization of complex tomographic data and supports the generation of high-quality figures for illustrative purposes.

### Camera paths and improvements for 3D animation

While ChimeraX already provides various options for creating publication-ready movies, the latest update of ArtiaX adds useful features for creating animations of cryoET data. With increased prevalence of samples prepared by FIB-SEM in recent years^12–14^ came the need for obliquely slicing data, which has been supported by ArtiaX since the first release. The updated plugin extends this to allow easy animation of obliquely oriented planes along any direction.

The 3D line primitive introduced above serves a dual purpose; it can be used both for modeling particle positions, but also to define camera flight paths during the recording of animations. Smooth paths are created by fitting cubic or higher degree splines to the selected points. The camera orientation is determined by its position along the path, and users can also explicitly set the camera’s rotation relative to the path.

### Image processing

ChimeraX provides functionality for low-pass filtering 3D images by means of gaussian filters. In the cryoEM space, bandpass filters other than traditional gaussians are frequently used (e.g a box filter with gaussian or raised cosine borders). Custom bandpass filters can be particularly useful when visualizing data with strong density gradients or curtaining artifacts. They can also be used to process noisy segmentations. ArtiaX now provides a user interface and commands to apply these filters.

Apart from this, ArtiaX now also allows computing moving averages (projection slices) along arbitrary directions of volumes. This method can be employed to increase the SNR of noisy tomograms and reduce the effects of the missing wedge distortion.

### Compatibility of particle lists with RELION-5 star files

CryoET data processing is a complex, multi-step procedure that often involves a variety of software tools. However, the use of different file formats across these tools can complicate the workflow, requiring file format conversions to transition between software packages.

To streamline the use of particle lists picked in ArtiaX across various software platforms, the plugin supports exporting files in multiple formats. This capability has now been extended to include saving particle lists in the new RELION-5^15^ .star file format, saving coordinates as centered coordinates in Angstroms and angle priors for particles with preorientations. ArtiaX also allows reading of particle star files generated after refinement or 3D classification jobs for assessment of refined orientations and positions of particles (Figure 3). A widget allows input of required fields such as the tomogram size and pixel size which are not contained in the header of the particle file but are needed for correct particle representation. If particle lists are modified and saved in the RELION5 format, the angle information is written out in the fields (rlnAngle<Rot/Tilt/Psi>) and (rlnTomoSubtomogram<Rot/Tilt/Psi>) conforming to a file created by a RELION5 Picks job, which can then be considered by relion_refine. A new particle list should be created for each tomogram separately, and entries should later be concatenated into one star file.

**Figure 3.**
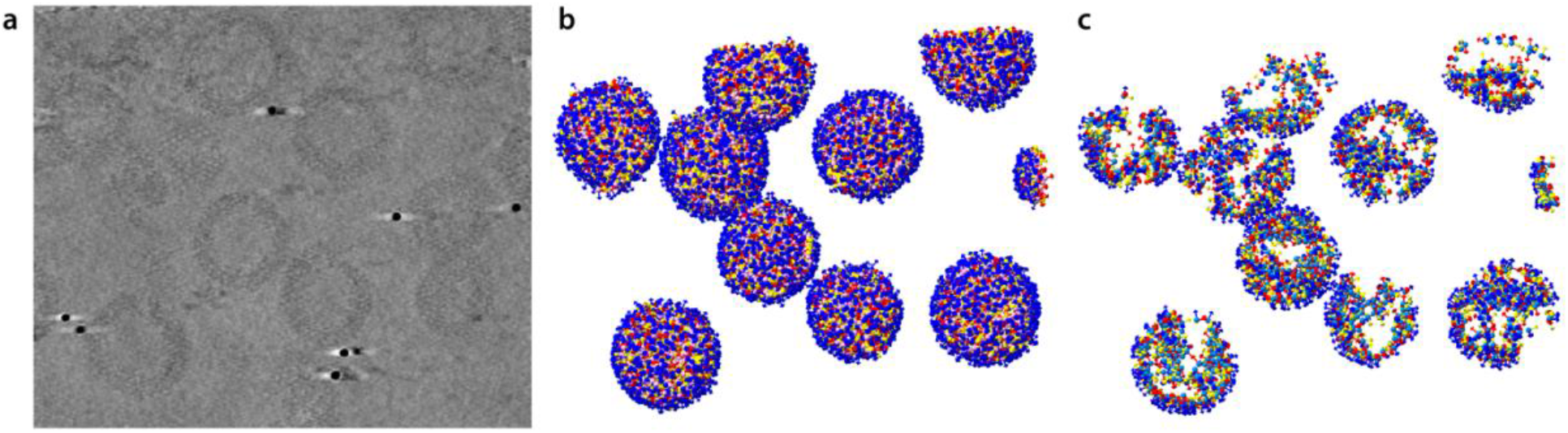
Compatibility of particle lists with RELION-5 star files. **(a)** Orthoslice through the tomogram TS_45 of the RELION5 tutorial dataset from EMPIAR10164. **(b)** Particle positions displayed from run_data.star after refinement (job12 precalculated results). **(c)** Particle positions displayed from class 5 of run_it025_data.star after 3D classification (job17 precalculated results).

Particle files can be loaded and saved without the need for loading a tomogram which allows rapid conversion between formats. Usage of the ChimeraX command line for batch processing is also enabled with the syntax for importing and saving to be found under “help artiax open/save”. Further documentation and tutorials are available under “help artiax user guide”.

## Discussion

ChimeraX stands out as a widely embraced tool for visualizing multiscale data and molecular models in the structural biology community^2,6^. Our plugin extends its capabilities by incorporating essential features tailored for extracting information from 3D image datasets, with a particular focus on meeting the specific requirements of electron tomography. This objective is accomplished without compromising user responsiveness, ensuring accessibility for all, and maintaining compatibility with the built-in ChimeraX functionality.

Looking ahead, a key future objective is the seamless integration of prior knowledge through easily deployable geometric or mesh-based models within a virtual reality (VR) context. Ongoing development efforts encompass manual and automatic tomogram segmentation^8,16^, tomogram denoising^17–19^, and pattern recognition within tomograms^17,20^. In forthcoming releases, we aim to expand ChimeraX functionality to incorporate tools addressing these challenges. Leveraging an open-source, extensible platform, we also adopt contributions from fellow researchers in the field.

### Software statement

All software is open source and can be download either from the ChimeraX toolshed or from GitHub. All the code is written in scientific python. This plugin uses several open-source python packages, including superqt for GUI elements, starfile for reading and writing RELION star files and pydantic for implementation of data models for particles.

## Author contributions

UHE, PR and GA developed python code. MW provided algorithms. DM and MPS tested software and implemented software manageability/availability. MPS and ASF designed research. UHE, DM, MPS, and ASF wrote the manuscript.

## Acknowledgments

We cordially thank Bogdan Toader for testing of ArtiaX for RELION5 and for his valuable input and advice for python coding. We also cordially thank Rangana Warshamanage for advice regarding handling the RELION5 format in python. UHE, PR, MW, DM & ASF were funded by Research Training Group iMOL (GRK 2566/1 and FR 1653/14-1).

## Declaration of interests

The authors declare that they have no competing interests.

## Notes

https://github.com/FrangakisLab/ArtiaX/

https://www.youtube.com/@artiax43

### Competing Interest Statement

The authors have declared no competing interest.

### Summary of Updates

Additions to the section on compatibility of ArtiaX with RELION-5; author list updated.

## References

1. Pruggnaller, S., Mayr, M. & Frangakis, A. S. A visualization and segmentation toolbox for electron microscopy. J. Struct. Biol. 164, 161–165 (2008).

2. Goddard, T. D. et al. UCSF ChimeraX: Meeting modern challenges in visualization and analysis. Protein Sci. Publ. Protein Soc. 27, 14–25 (2018).

3. Sofroniew, N. et al. napari: a multi-dimensional image viewer for Python. Zenodo 10.5281/zenodo.13309520 (2024).

4. Dragonfly 2024.1 [Computer software]. Comet Technologies Canada Inc., Montreal, Canada; software available at https://www.theobjects.com/dragonfly.

5. Ermel, U. H., Arghittu, S. M. & Frangakis, A. S. ArtiaX: An electron tomography toolbox for the interactive handling of sub-tomograms in UCSF ChimeraX. Protein Sci. Publ. Protein Soc. 31, e4472 (2022).

6. Pettersen, E. F. et al. UCSF ChimeraX: Structure visualization for researchers, educators, and developers. Protein Sci. Publ. Protein Soc. 30, 70–82 (2021).

7. Wagner, T. et al. SPHIRE-crYOLO is a fast and accurate fully automated particle picker for cryo-EM. Commun. Biol. 2, 1–13 (2019).

8. de Teresa-Trueba, I. et al. Convolutional networks for supervised mining of molecular patterns within cellular context. Nat. Methods 20, 284–294 (2023).

9. Eibauer, M. et al. Vimentin filaments integrate low-complexity domains in a complex helical structure. Nat. Struct. Mol. Biol. 31, 939–949 (2024).

10. Sikora, M. et al. Desmosome architecture derived from molecular dynamics simulations and cryo-electron tomography. Proc. Natl. Acad. Sci. U. S. A. 117, 27132– 27140 (2020).

11. Vizarraga, D. et al. Immunodominant proteins P1 and P40/P90 from human pathogen Mycoplasma pneumoniae. Nat. Commun. 11, 5188 (2020).

12. Berger, C. et al. Cryo-electron tomography on focused ion beam lamellae transforms structural cell biology. Nat. Methods 20, 499–511 (2023).

13. Noble, A. J. & de Marco, A. Cryo-focused ion beam for in situ structural biology: State of the art, challenges, and perspectives. Curr. Opin. Struct. Biol. 87, 102864 (2024).

14. Kelley, K. et al. Waffle Method: A general and flexible approach for improving throughput in FIB-milling. Nat. Commun. 13, 1857 (2022).

15. Burt, A. et al. An image processing pipeline for electron cryo-tomography in RELION-5. FEBS Open Bio (2024).

16. Chen, M. et al. Convolutional neural networks for automated annotation of cellular cryo-electron tomograms. Nat. Methods 14, 983–985 (2017).

17. Tegunov, D., Xue, L., Dienemann, C., Cramer, P. & Mahamid, J. Multi-particle cryo-EM refinement with M visualizes ribosome-antibiotic complex at 3.5 Å in cells. Nat. Methods 18, 186–193 (2021).

18. Liu, Y.-T. et al. Isotropic reconstruction for electron tomography with deep learning. Nat. Commun. 13, 6482 (2022).

19. Buchholz, T.-O., Jordan, M., Pigino, G. & Jug, F. Cryo-CARE: Content-Aware Image Restoration for Cryo-Transmission Electron Microscopy Data. Preprint at 10.48550/arXiv.1810.05420 (2018).

20. Wan, W., Khavnekar, S. & Wagner, J. STOPGAP : an open-source package for template matching, subtomogram alignment and classification. Acta Crystallogr. Sect. Struct. Biol. 80, 336–349 (2024).

